# Challenging the boundaries of the physical self: purely distal cues in the environment impact body ownership

**DOI:** 10.1101/672139

**Authors:** Klaudia Grechuta, Javier De La Torre, Belén Rubio Ballester, Paul F.M.J. Verschure

## Abstract

The unique ability to identify one’s own body and experience it as one’s own is fundamental in goal-oriented behavior and survival. However, the mechanisms underlying the so-called body ownership are yet not fully understood. The plasticity of body ownership has been studied using two experimental methods or their variations. Specifically, the Rubber Hand Illusion (RHI), where the tactile stimuli are externally generated, or the *moving* RHI which implies self-initiated movements. Grounded in these paradigms, evidence has demonstrated that body ownership is a product of bottom-up reception of self- and externally-generated multisensory information and top-down comparison between the predicted and the actual sensory stimuli. Crucially, provided the design of the current paradigms, where one of the manipulated cues always involves the processing of a proximal modality sensing the body or its surface (e.g., touch), the contribution of sensory signals which pertain to the environment remain elusive. Here we propose that, as any robust percept, body ownership depends on the integration and prediction of all the sensory stimuli, and therefore it will depend on the consistency of purely distal sensory signals pertaining to the environment. To test our hypothesis, we create an embodied goal-oriented task and manipulate the predictability of the surrounding environment by changing the congruency of purely distal multisensory cues while preserving bodily and action-driven signals entirely predictable. Our results empirically reveal that the way we represent our body is contingent upon all the sensory stimuli including purely distal and action-independent signals which pertain to the environment.

## Introduction

The sense of body ownership, which allows us to determine the boundaries between the own physical self and the external world, and therefore the source of a given sensation, is fundamental in adaptive goal-oriented behavior and survival [11, 16, 71, 74]. Indeed, during the last three decades, scientists have increasingly questioned both the behavioral and neural mechanisms driving the emergence and experience of body ownership as well as its flexibility [2, 5, 6, 20, 23, 66]. Together, the results support the notion that the way we perceive our body strongly relies on an interplay between (1) bottom-up reception, combination, and integration of self-generated (reafferent) and externally-generated (exafferent) information from multiple sensory sources, and (2) top-down comparison between the expected and the actual sensory stimuli [2, 5, 6, 68]. At the empirical level, the principles underlying bodily representation (in healthy subjects) have been studied using bodily illusions [13]. A well-established experimental paradigm is the Rubber Hand Illusion (RHI) which is used to induce ownership over a fake rubber hand by manipulating the congruency of externally-delivered tactile cues [6]. Another standard method for inducing ownership is the so-called *moving* Rubber Hand Illusion (mRHI). Here, the visual feedback of self-initiated arm or finger movements is either synchronized with the actual trajectory or not, thus biasing the feeling of ownership over the virtual body [18, 38, 58, 70]. Crucially, both in the RHI and mRHI or their variations, one of the sensory signals manipulated to induce the experience of ownership always involves the processing of a proximal modality such as touch or proprioception [6, 58]. Consequently, the current understanding of the mechanisms driving body ownership is constrained to the study of sensory cues which pertain to the body or the sensory consequences of its self-initiated movements within the peripersonal space (PPS) [8, 9, 37, 48, 49, 54, 55]. Indeed, only recently it has been shown that in the context of a goal-oriented action, such as hitting the puck in Air Hockey, body ownership is also modulated by purely distal (i.e., auditory) cues which pertain to the task [33]. Interestingly, however, the contribution of purely distal signals which are action-independent and which pertain exclusively to the environment is still not fully understood.

With seemingly no effort, we generate unambiguous interpretations about the self and the environment and determine the boundaries between them [16]. To do so, our brain uses multiple sources of sensory information processed by different modalities (i.e., vision, touch, audition) [22]. As such, any robust percept including the sense of ownership requires merging of this information which continuously occurs within and outside of the body that is in the environment [22]. For instance, we simultaneously receive and integrate inputs informing about the location and position of our limbs as well as those informing about the location and position of objects in the scene. Until now, however, the experimental evidence about the multisensory representation of the body and the necessary and sufficient conditions for the experience of its ownership is grounded exclusively in the study of proximal cues [5, 6, 23, 31, 42, 66]. For instance, the seminal experiment of Botvinick & Cohen [6] and later many others (for review see [66]) who used RHI as a method to manipulate body ownership, proposed that the self-attribution of the rubber hand arises reactively as a result of bottom-up processes of combination and integration of information from visual and tactile modalities. Hence, initially, the illusion of owning the fake hand was interpreted as a passive perceptual state whose strength was correlated with temporal discrepancies between seen and felt sensory stimuli (both necessary and sufficient). In the light of recent findings, however, this traditional view on body ownership as resulting purely from perceptual correlations does not seem sufficient [66]. In particular, it has been widely accepted that despite its flexibility, in the context of externally-generated sensory cues (e.g., tactile strokes as in the RHI), body ownership somewhat requires physical, anatomical, postural and spatial congruency of the real (felt) and fake (viewed) hands [14, 25, 35, 45, 48, 68]. These findings strongly suggest that the interpretation of the ‘novel’ sensory evidence and possible incorporation of the rubber hand into the representation of the body is constrained by top-down prior knowledge driven by experience [2, 5, 44, 68]. In particular, the perception of ownership seems to rely on an internal model of the body which relates the physical aspects of the perceived rubber hand to the inputs received through a history of sensorimotor interactions of an agent with the world [2, 66]. Interestingly, this hypothesis is consistent with the general framework which proposes that perception is controlled by top-down processes allowing to create predictions about the forthcoming sensory events based on previous experience and generalized knowledge [21, 27, 40]. As such, it is an active process in which all acquired sensory information is continuously compared against experience-driven internal models of the self and the environment [7, 10, 23, 47, 53].

Grounded in the framework of active perception, Ferri and colleagues [23, 24] studied whether body ownership can be modulated by a pure expectation of exafferent tactile feedback in the absence of actual physical touch of either the fake or the real body-parts. Interestingly, their experiment revealed that a mere expectation of an upcoming sensory event, predicted by an anticipatory response in multisensory parietal cortices, is indeed sufficient to induce the experience of ownership over the rubber hand, measured subjectively and objectively. However, the tactile stimulation is not necessary [23, 24] (see also [62]). This result emphasizes the predictive processing in the emergence of the sense of body ownership challenging the traditionally defined boundaries of an embodied self. In the present study, we extend this hypothesis and propose that, as any coherent percept, body ownership is a result of bottom-up integration and top-down prediction of all the sensory stimuli processed by proximal and distal modalities including those which pertain purely to the environment. Hence, we propose that body ownership will depend on the consistency of distal sensory signals which occur in the environment even if they are independent of self-initiated actions. To test this hypothesis, we create an embodied goal-oriented task using virtual reality and manipulate the predictability of the surrounding environment by changing its rules while preserving bodily and action-driven signals fully predictable. We predict that body ownership of a virtual avatar will be negatively influenced in the condition where purely external sensory signals underlying the statistical structure of the environment are not predictable.

## Materials and Methods

### Participants

Twenty-four healthy naive students from Universitat Politècnica de Catalunya provided their consent and participated in the study: 12 males (Mean ± SD 23,25 ± 2,37 years-old) and 12 females (Mean ±SD 22,16 ± 2,12 years-old). The sample size was chosen based on previous studies [41, 61]. All the subjects were right-handed (handedness assessed using the Edinburgh Handedness Inventory [52]), had normal hearing and normal or corrected-to-normal vision. Additionally, none of the participants reported stereoblindness. Each participant was pseudorandomly assigned to one of the two experimental conditions, following a between-subjects design. Similar to [3], this experimental method was chosen to prevent habituation to the ownership measures and sensory manipulations which could bias the physiological responses in the subsequent blocks. The groups were balanced with respect to age, gender and previous experience with virtual reality. We ensured that none of the participants had previously participated in a body-ownership study, and informed the subjects that they were free to withdraw from the experiment at any time.

### Experimental setup, procedures, and protocol

#### Experimental Setup

The present experimental setup consisted of a personal computer, head mounted display (HTC Vive, www.vive.com), motion detection input device (Kinect, Microsoft, Seattle, USA), and active noise control headphones (Beats Electronics LLC, California, USA) which served to ensure isolation from external sounds (Figure 1B). Similar to previous experiments [31, 58], here we used the virtual reality method as a tool to investigate the modulation of body ownership [57]. The protocol was integrated within the Rehabilitation Gaming System (RGS) [32], the Virtual Environment (VE) was designed using SketchUp (Trimble Inc., California, USA) (Figure 1C), and the software was developed in Unity3D (Unity Technologies SF, Copenhagen, Denmark). During the experiment, subjects sat at a cut-out table with their arms resting (Figure 1B). The movements of the arms were continuously tracked and mapped onto the avatar’s arms enabling interaction with the VE.

**Figure 1.**
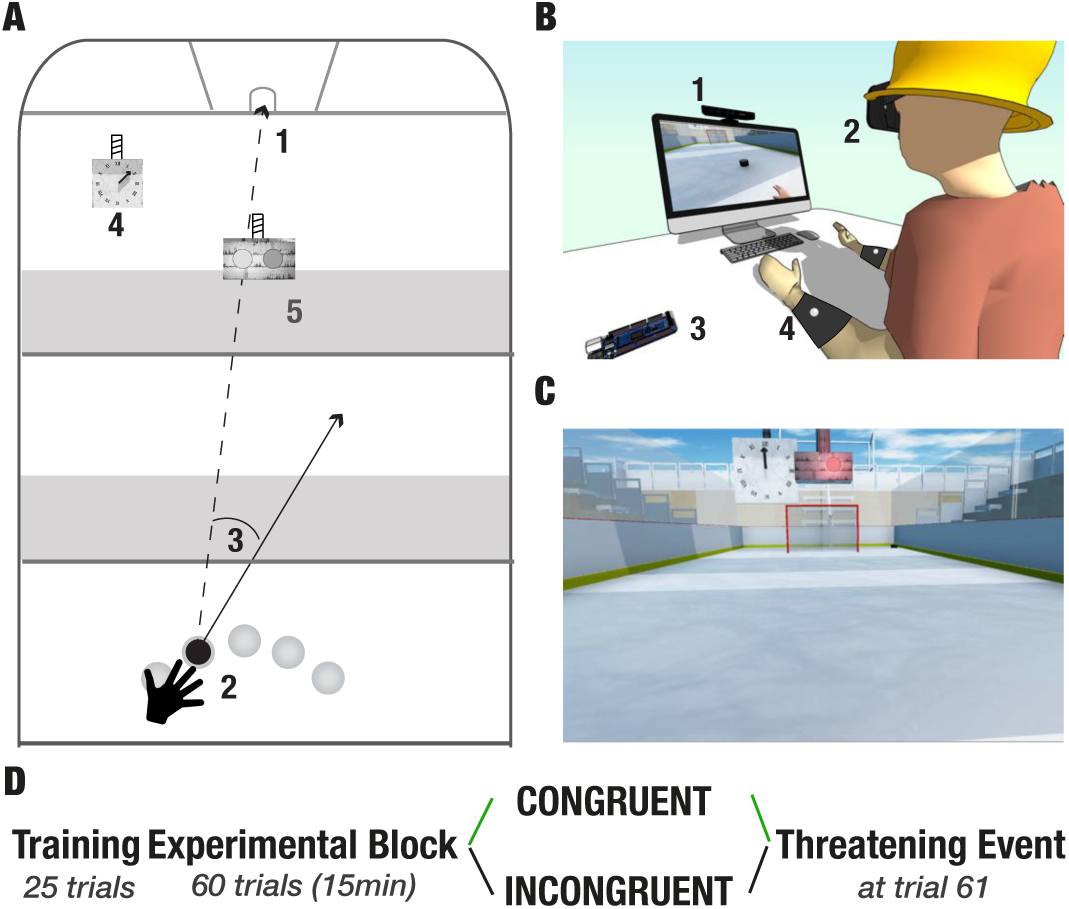
Experimental setup and protocol. **(A) Schematic bird view of the virtual environment.** 1-goal; 2-five starting positions; 3-an example of a directional error; 4-the Go Signal (GS); 5-a clock. **(B) Experimental setup.** 1-motion tracking device (Kinect); 2-Head Mounted Display (HMD, HTC-Vive Headset); 3-Arduino & E-Health; 4-Bracelets with reflecting markers for the tracking. **(C) Virtual scene.** The first person perspective view of the scene. **(D) Experimental protocol.** All participants go through the *Training Block* (25 trials). Then they are randomly split into two experimental conditions: congruent (“C”, green arrow) and incongruent (“I”, black arrow) and they undergo the *Experimental Block*. At the end of the experiment, all the participants experience the *Threatening Event*.

#### Virtual Air Hockey Task

The objective of each session was to complete a goal-oriented motor task that consisted of hitting a virtual puck into a goal, as accurately as possible (Figure 1A). Prior to the experiment, subjects received instructions to place their hands in square-shaped Go Areas (GA; one for the left and one for the right hand) at the beginning of every trial. The trial started, and the puck appeared only when the system detected that both hands are in the GAs. Importantly, although both hands were mapped and rendered in the virtual scene, the task was to be completed using the right arm exclusively. To counteract repetitive movement-patterns and prevent habituation the puck was spawned pseudorandomly in one of the five Starting Positions (SP) distributed evenly within the right-hand workspace (Figure 1A2). The game was designed such that the puck did not bounce against the walls. Thus a trial consisted of one hit only which could end in either a success (i.e., the puck enters the goal) or a failure (i.e., the puck hits one of the walls). Both events were immediately indicated by auditory feedback in the form of semantically corresponding sound, positive or negative respectively. Pertaining to the task, this feedback was always congruent and fully predictable. The puck was visible throughout the experiment.

#### Protocol and Sensory Manipulations

The experiment consisted of three parts including (1) Training Block (TB, 25 trials), (2) Experimental Block (EB, 60 trials), and (3) a Threatening Event (TE) (Figure 1D). TB and TE were the same for all the participants. TB, in particular, allowed familiarization with the virtual environment as well as with the dynamics of the game, while TE served to record autonomous, physiological responses to an unexpected threat as an objective measure of body ownership [3]. In the EB, subjects were randomly split into two groups: Congruent “C” (green), and Incongruent “I” (black) (Figure 1D). To investigate whether sensory cues which pertain to the environment influence body ownership in the experimental condition (“I”), we manipulated the congruency, and therefore the predictability, of visual and auditory action-independent cues from the virtual scene.

The scene consisted of (1) the virtual arms, (2) an Air Hockey field, (3) the puck, (4) a goal, (5) benches for the audience, (6) the Go Signal (GS), and (7) a clock (Figure 1). The virtual solar time was indicated by the position of the sun in the sky (i.e., visual cue), while the virtual space (setting) was signaled by the background sound representative for a given place (i.e., auditory cue; e.g., the sound of the air-hockey field) (Figure 1C). The default scene was set at midday (setting:time) on a hockey field (setting:location). Both in the Training Block (identical for both conditions) and the Experimental Block of the Congruent condition “C” all the scene components mentioned above, as well as the temporal and spatial settings, remained fixed such that their behavior was always fully predictable. Moreover, in all conditions, all the auditory and visual signals relevant to the body within the peripersonal space and to the task (i.e., Air Hockey field, the puck, the goal, and the Go Signal, the trajectory of the puck, outcome of the action) were always congruent and fully predictable. Crucially, in the Experimental Block of the Incongruent condition (“I”), the default behavior of the scene components and the temporal and spatial settings randomly changed. In particular, we manipulated: (1) spatial orientation of the benches by rotating them on the z-axis, (2) spatial orientation of the clock by modulating the velocity and the direction of the arrows indicating the time, (3) virtual solar time by altering the position of the sun in the sky or changing its characteristics to nighttime, and (4) virtual space by altering the background sound (i.e., sounds representative for a concert, cinema). Importantly, to ensure that the sensory manipulations in “I” impact exclusively the perception of the environment and not the action, they were always introduced between the end of a trial (the puck enters the goal or hits one of the walls) and the beginning of the consecutive one. Specifically, they were triggered at a random time within a 2 seconds time window after the end of each trial. The incongruencies were introduced gradually, and they were pseudorandomly distributed such that the participants could not attribute action-driven causality to their emergence.

### Measures

#### Self-reports

In virtual environments, the sense of presence refers to the subjective experience of ‘being there’, despite the physical distance. In particular, when a user does not perceive the influence of technology during a virtual reality-based experience [57, 72]. To ensure that the participants in both groups felt equally immersed within the proposed environment, we asked them to complete a presence questionnaire at the end of the experiment by assessing each of the items on a 9-point Likert Scale (see Table2 for the full list of items). Furthermore, to evaluate the subjective experience of body ownership and agency, we administered a 12 item questionnaire (Table 1) adapted from previous studies [38, 46]. There were six questions per domain, three of which served as controls. Participants answered each statement on a 7-point Likert Scale ranging from ‘-3’: being in strong disagreement to ‘3’: being in strong agreement. To counteract order effects, the sequence of questions was randomized.

**Table 1.** The questionnaire, consisting of 12 statements divided into four different categories.

| Category | Question |
| --- | --- |
| Ownership | I felt as if I was looking at my own hand |
|  | I felt as if the virtual hand was part of my body |
|  | I felt the virtual hand was my hand |
| Ownership Control | It seemed as if I had more than one left hand |
|  | It appeared as if the virtual hand were drifting towards my real hand |
|  | It felt as if I had no longer a left hand, as if my left hand had disappeared |
| Agency | The virtual hand moved just like I wanted it to, as if it was obeying my will |
|  | I felt as if I was controlling the movements of the virtual hand |
|  | I felt as if I was causing the movement I saw, and the control questions were |
| Agency Control | I felt as if the virtual hand was controlling my will |
|  | I felt as if the virtual hand was controlling my movements |
|  | I could sense the movement from somewhere between my real and virtual hand |

#### Galvanic Skin Response (GSR)

GSR is a physiological measure of the autonomous nervous system, which increases as a reaction to an arousing stimulus. Similar to other studies [3], here, we used GSR as a proxy for the ownership illusion. In particular, at the end of the Experimental Block in each condition, we measured GSR responses to an unexpected threat (i.e., a knife falling to stab the right virtual hand). To prevent movement-driven muscular artifacts, we recorded GSR from the left hand which did not move during the experiment. The signal was recorded using Arduino e-Health board at a sampling rate of 33Hz from two flat reversible silver/silver chloride (Ag–AgCl) electrodes which were attached to the middle and index fingers, respectively. We stored the GSR during the whole experiment interval for each participant. The data were preprocessed to extract phasic components from tonic activity based on Continuous Decomposition Analysis (CDA) [4] as implemented in the Ledalab software (Leipzig, Germany). For the analysis, we computed the number of phasic skin conductance responses (nSCR) above 0.01mS 1 second following the threatening event and compared it between the two conditions.

#### Hand Withdrawal (HW)

We collected kinematic movement data from the Kinect for each participant throughout the experiment. All the data from the system was recorded at 33Hz. To quantify the execution of instinctive defensive movements such as hand withdrawal in response to the unexpected threatening event (i.e., the virtual knife stabbing the virtual hand) [30], we computed the velocity of displacement of the right virtual hand as a difference in the cumulative sum of the X (forward and backward) and Y (up and down) positions at every time step. The results were compared between the two conditions. Due to possibly stronger assimilation of the virtual hand to the representation of the body, we expected higher velocity of movement in “C” than in “I”.

#### Performance measures

To evaluate performance, we measured scores, angular errors as well as reaction and response times, all stored by the system. Scores were calculated as the percentage of successful trials, namely, the times when the puck entered the gate (Figure 1A1). An angular error was computed as the difference between the actual direction vector and a straight line between the starting position of the puck and the middle of the goal (desired trajectory, Figure 1A3). Reaction times were the time intervals between the apparition of the puck and the moment of ‘leaving’ the starting position to hit it, while the response times were the time intervals between the apparition of the puck and the moment of its collision with the hand.

The statistical analysis followed nonparametric methods. Hence, we used the Mann–Whitney U test for between groups analyses and the Wilcoxon signed-rank test for within groups comparisons. The data were analyzed using Python3.6.4 (http://www.python.org) and Matlab (Mathworks, Natick, USA).

## Results

To determine whether body ownership depends on the consistency of purely external cues which pertain to the environment, we developed an embodied goal-oriented task and manipulated the congruency, and therefore the predictability of action-independent sensory cues outside of the peripersonal space while preserving bodily and action-driven cues fully predictable. For each session and each participant in both conditions, we used Galvanic Skin Response (GSR), Hand Withdrawal (HW), and self-reports to quantify body ownership objectively, behaviorally and subjectively. Furthermore, we stored performance scores, angular errors as well as response and reaction times as measures of performance. Finally, to ensure that the participants were immersed in the proposed virtual environment and perceived the sense of agency over the virtual limbs, we administered *presence* (adapted from [72]) and *agency* questionnaires (adapted from [46]).

### Presence and agency

First, we assessed the perceived experience of presence and agency. The analysis revealed that in both conditions participants felt present in the proposed virtual environment (“C”: *µ* = 1.6, *std* = 1.56 and “I”: *µ* = 1.51, *std* = 1.8) (Figure 2). Crucially, we found no differences between the groups in the self-reported scores (*p* = 0.47). Table2 presents individual questionnaire items as well as the results of between-group analyses for the associated questions. None of the comparisons yielded a statistically significant difference. We further report no difference in the experienced sense of agency between “C” (*µ* = 1.19, *std* = 1.24) and “I” (*µ* = 1.3, *std* = 1.3) (*p* = 0.08) (Figure 2), and the groups did not differ in the agency control questions (*p* = 0.1).

**Figure 2.**
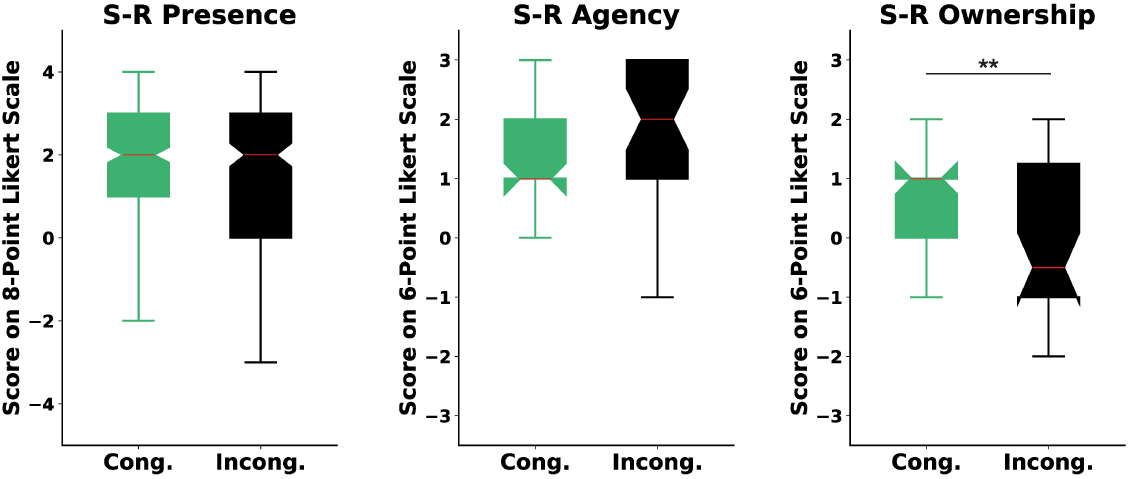
Self Reports. **Left.** Self-reported experience of presence. Y-axis: Responses on the 9-point Likert scale ranging from ‘-4’ (strongly disagree) to ‘4’ (strongly agree). Scores above ‘0’ indicate a feeling of presence. **Middle.** Self-reported experience of agency. Y-axis: Responses on the 7-point Likert scale ranging from ‘-3’ (strongly disagree) to ‘3’ (strongly agree). Scores above ‘0’ indicate a feeling of agency. **Right.** Self-reported experience of ownership. Y-axis: Responses on the 7-point Likert scale ranging from ‘-3’ (strongly disagree) to ‘3’ (strongly agree). Scores above ‘0’ indicate a feeling of body ownership.

### Performance

Our results revealed that the normalized performance-scores (i.e., the proportion of successful trials) did not differ between the Congruent (*µ* = 0.6, *std* = 0.17) and the Incongruent (*µ* = 0.59, *std* = 0.18) conditions (*p* = 0.5). We further analyzed the angular error as a proxy to performance, stored throughout the experiment. We report that while both groups significantly improved during the Training Block (TB), in the Experimental Block (EB) the errors stabilized (Figure 3). Specifically, there was a statistically significant difference between the early and late trials in both “C” (early trials: *µ* = 12.63, *std* = 5.25 and late trials: *µ* = 6.11, *std* = 3.76; *p* = 0.002) and “I” (early trials: *µ* = 14.03, *std* = 6.9 and late trials: *µ* = 7.53, *std* = 2.48; *p* = 0.004) in the Training Block. We found, however, no within-group differences for the early and late trials in the Experimental Block: “C” (early trials: *µ* = 8.05, *std* = 3.81 and late trials: *µ* = 7.14, *std* = 4.87; *p* = 0.53) and “I” (early trials: *µ* = 7.1, *std* = 3.85 and late trials: *µ* = 7.26, *std* = 3.94; *p* = 0.48). Furthermore, the Mann–Whitney U test yielded no differences in performance (angular error) between “C” and “I” in neither the TB (*p* = 0.15) nor the EB (*N* = 10, *p* = 0.15) (Figure 3) demonstrating that, overall, the conditions did not differ with respect to performance. Finally, our analysis showed no statistically significant differences in either response (“C”: *µ* = 2.35, *std* = 0.78 and “I”: *µ* = 2.43, *std* = 0.67; *p* = 0.1) or reaction times (“C”: *µ* = 1.01, *std* = 1.41 and “I”: *µ* = 1.63, *std* = 1.14; *p* = 0.3) between the two conditions during the Experimental Block. Hence both groups took the same time to initiate the movement and hit the puck, further suggesting that the proposed manipulation of action-independent sensory signals in “I” did not alter or bias performance.

**Figure 3.**
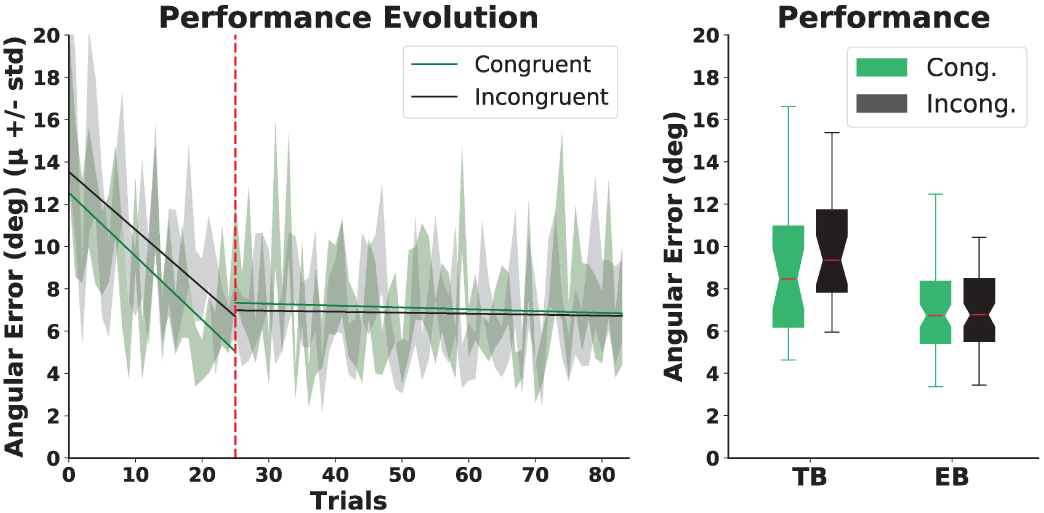
Motor performance. **Left.** The Evolution of Angular Errors. The dashed red line indicates the end of the Training Block (TB) and the beginning of the Experimental Block (EB). The solid lines represent linear regression models for the angular errors in each condition in the Training Block and Experimental Blocks, before or after the dashed red line respectively. **Right** Total Angular Errors. Boxplots represent angular errors for the two conditions in the training and Experimental Blocks respectively. No differences were found between the groups.

### Body Ownership

The analysis revealed a statistical difference in the self-reported experience of body ownership between the two conditions (*p* = 0.04) such that the scores were significantly higher in “C” (*µ* = 0.6, *std* = 0.9) than in “I” (*µ* = 0.05, *std* = 1.45) (Figure 2). Importantly, we found no differences in the control questions between the groups (*p* = 0.1). To further explore the effects of the congruency of purely external cues on body ownership, we computed post-threatening GSR responses for every individual in both groups (Figure 4). The signal per each participant was normalized by subtracting the mean signal from 10 seconds prior to the stimulus onset.As expected and in line with the literature, the GSR signal increased in both groups. Specifically, the threatening event triggered a significant increase in the number of galvanic skin responses in “C” (pre-TE: *µ* = 1, *std* = 0.577, post-TE: *µ* = 2.58, *std* = 1.11, *p* = 0.0008) and in “I” (pre-TE: *µ* = 1.16, *std* = 0.55, post-TE: *µ* = 2.33, *std* = 1.64, *p* = 0.03). Crucially, however, we found a statistically significant difference between the groups in the numbers of activations (*p* = 0.003) such that the number was significantly higher in “C” (*µ* = 2.58, *std* = 1.11) than in “I” (*µ* = 1.833, *std* = 0.98) (Figure 4).

**Figure 4.**
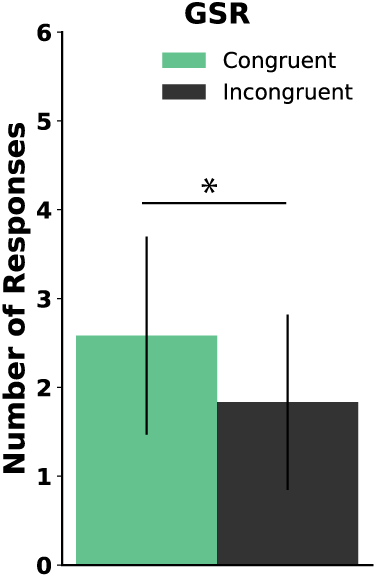
GSR results. The plot represents the difference between the groups in the number of galvanic skin responses (nGSR) post stabbing event.

Finally, we observed that in the congruent condition participants exhibited faster velocity of the right virtual hand displacement post threatening event (Hand Withdrawal, Figure 5). In particular, the statistical analysis revealed that the difference between “I” and “C” in the cumulative sum of the X and Y position over time reached statistical significance at second 4 post threatening event (Figure 5).

**Figure 5.**
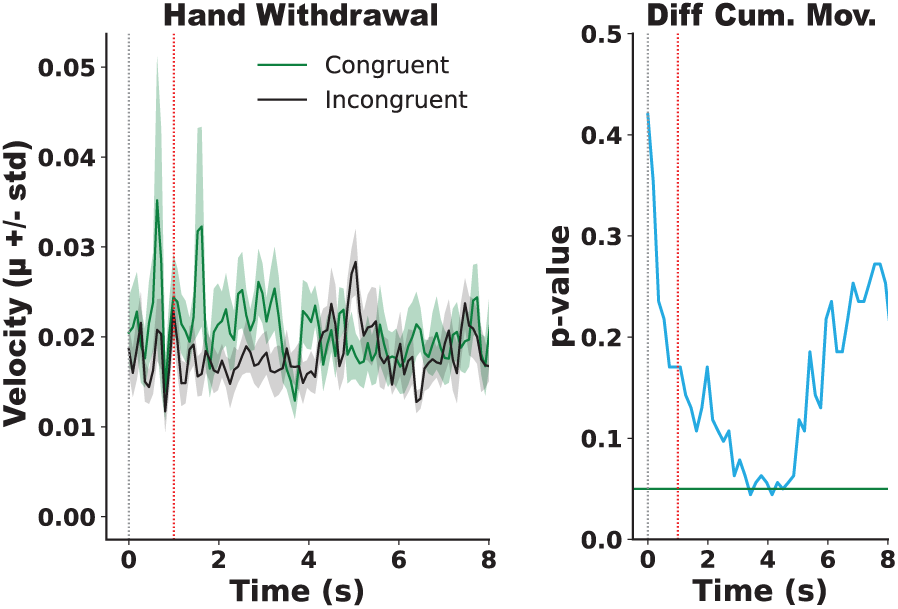
Hand Withdrawal results. **Left.** Changes in Movement Velocity Post-Threatening Event. The graph represents the evolution of changes in X and Y positions of the right virtual hand following the threatening event for each group over time. The data on the X-axis was recorded at 33Hz, and it is presented in an overlapping window of 66ms. **Right.** Changes in Velocity Between the groups. The blue line represents *p values* obtained from Mann-Whitney U tests performed on the cumulative sum of the movement velocity. The values corresponding to each time step on the X-axis are shown in overlapping windows of 150-ms. In B, the green horizontal line indicates 0.05 significance threshold. In both graphs, the dashed gray line indicates the time when the knife appeared (*time* = 0) while the dashed red line shows the time when the knife stabbed the hand (*time* = 1).

## Discussion

The unique ability to recognize one’s own body, experience it as our own, and localize it in space lies in the continuous processing of self- (i.e., reafferent) and externally-generated (i.e., exafferent) multisensory information arising from sensorimotor interactions of an agent within the environment [11, 16, 39, 74]. Vast evidence has now demonstrated that this processing comprises bottom-up reception and top-down prediction of sensory stimuli which pertain to the body and occur within the peripersonal space [6, 19, 24, 50, 58, 59]. The present study extends prior findings by showing that the plasticity of body ownership also depends on the consistency of body- and action-independent sensory cues which pertain to the environment and are*-* processed by purely distal modalities (i.e., visual and auditory). Thus, for the first time, we empirically reveal that the way we represent our body is contingent upon all the sensory stimuli including signals occurring outside the peripersonal space. We interpret our results from the perspective of active perception and propose that, similar to any robust percept, body ownership depends on the consistency of the internal models of not only the body or the consequences of its actions but also the model of the surrounding environment [16, 21, 22, 51].

A large body of evidence demonstrates that the experience of ownership emerges actively through dynamic comparisons between integrated and predicted multisensory signals [2, 23, 33, 60, 65, 66]. The influence of top-down processing [65] on the sense of ownership is supported in the contexts of classical Rubber Hand Illusion (RHI) where the stimuli are externally generated [6]. Here, self-attribution of the rubber hand arises actively as a consequence of the minimization of prediction errors resulting from multisensory conflicts during the synchronous stroking of the real and fake body-parts (i.e., visuotactile) [2]. Similar, a failure to experience ownership over a noncorporeal object [67] or a rubber hand located in an implausible position [14] can also be interpreted as a consequence of predictive matching of the sensory inflow and the experience-driven internal models of the body [2]. Moreover, it has been recently demonstrated that ownership can be induced by a pure expectation of correlated sensory input [23, 24, 62].

The contribution of top-down processing in the emergence of body ownership is further supported in the context of self-generated cues such as in the *moving* Rubber Hand Illusion (RHI) [18, 57]. Here, it has been proposed that the location of different body-parts is estimated by the Central Nervous System (CNS) via a Forward Model (FM) or a Corollary Discharge (CD) which generates predictions about the sensory consequences of movements and compares them with the corresponding sensory feedback [51, 63, 75]. Those predictions are carried out by the so-called efference copy employing all the sensory signals relevant to the body and the goal of the task (i.e., task-relevant) [34, 64, 73]. The Sensory Prediction Errors (SPE) from multiple sensory sources, which reflect the discrepancies between the expected and the actual sensory stimuli, inform the brain about the current state of the environment and the body, shaping the experience of ownership. For instance, when the visual feedback of the virtual hand does not match the expected one (e.g., the hand follows a different trajectory than the executed one) the SPEs pertaining to the body increase resulting in a decreased experience of ownership over the virtual avatar [18, 58]. Interestingly, even purely distal auditory consequences of self-generated movements bias the experience of ownership provided that they are relevant to the task, thus informing about the magnitude of the error [33].

The evidence discussed above suggests that body ownership is compromised when the actual sensory signals violate the expected cues independently of whether they are externally-(RHI) or self-generated (mRHI). If body ownership depends on the matching between the predicted and the actual sensory stimuli, can it, in a similar way, be affected by prediction errors about the sensory signals pertaining to the environment? To answer this question, we designed a virtual reality-based paradigm where participants were to complete a motor task (Air Hockey) and manipulated the predictability of the purely external cues by randomly changing the rules of the environment. Thus, similar to the prediction errors which result from visuotactile matching and affect the internal (generative) model of the body [6], or those which result from visuomotor [58] or visuoauditory [33] matching and affect the internal (forward) model, here we experimentally induced prediction errors which result from visuoauditory matching and affect the internal (generative) model of the environment [12, 26]. We expected that if body ownership depends on the congruency of all the sensory stimuli, it will be impacted in the experimental block of the incongruent condition where the expectations about the model of the environment acquired during the training block are violated. Our findings establish that incongruencies in action-independent and task-irrelevant sensory cues, which inform about the statistical structure of the environment and are processed by purely distal modalities, modulate the experience of body ownership. In particular, we found that the congruent (as compared to incongruent) environment led to an enhanced experience of ownership over the virtual hand, as measured subjectively by a questionnaire (Figure 2), behaviorally using the hand withdrawal (Figure 5), and objectively through the galvanic skin responses (Figure 4). Crucially, however, there were no effects regarding the experience of presence (Table 2 and Figure 2) supporting that, despite the introduced manipulations, the environment was overall immersive [57].

**Table 2.** All items from presence questionnaire. P-values indicate the results of a between-group comparison all the items using the Mann–Whitney U test.

| question | p-value |
| --- | --- |
| Were you able to control events? | 0.179 |
| How responsive was the environment to actions you initiated | 0.257 |
| Naturality of the interaction | 0.232 |
| How much did visual aspects involve you? | 0.476 |
| How natural was the movement mechanism | 0.34 |
| How compelling was the sense of objects moving | 0.21 |
| How consistent were the experiences with real world ones | 0.384 |
| Were you able to anticipate consequences of your actions | 0.22 |
| Were you able to survey the environment using vision | 0.39 |
| How compelling was the sense of moving around | 0.36 |
| How closely were you able to examine objects | 0.22 |
| Were you able to examine objects from multiple viewpoints | 0.2 |
| How involved were you in the virtual reality experience | 0.18 |
| How much delay did you experience | 0.476 |
| How quickly did you adjust to the virtual reality experience | 0.4 |
| How proficient in interacting did you feel at the end | 0.439 |
| How much did visual aspects distract from the task | 0.38 |
| How much did the control devices interfere with performance | 0.373 |
| How well could you concentrate on the assigned tasks | 0.164 |
| How much did auditory aspects involve you | 0.064 |
| How well could you identify sounds | 0.474 |
| How well could you localize sounds | 0.21 |

We propose that the violation of expectations in the context of the proposed paradigm can be understood as a sudden increase of uncertainty in the internal model of the environment. According to biologically-constrained models of the neocortex, the link between these two components could be mediated by neuromodulators which signal uncertainty such as norepinephrine or acetylcholine [1]. Consequently, a significant enough violation of expectations influenced by the sensory manipulations in the incongruent condition would have a global effect on modulating uncertainty in a range of brain areas including those which underly the multisensory representation of the body (e.g., the right insula, posterior parietal and ventral premotor cortices) [15, 29, 69]. From the functional perspective, such sensory prediction errors would have the same impact on all predictive models inducing uncertainty in the model of the environment (i.e., a generative model) and the model of the body. In other words, the temporal modulation of global uncertainty would inevitably change the overall confidence in the internal model of the body as supported by our results (Figure 2, 4, 5). The present outcomes are further consistent with the Bayesian causal inference and could be interpreted in terms of likelihood according to which if the environment is likely *I* am likely too, and vice-versa [2, 17, 43, 56].

An interesting question which one could raise, however, is how persistent is this effect? Is it transient? It has been demonstrated that one way to minimize prediction errors is to update the current model, in our case, the model of the environment, to accommodate the unexpected sensory signals [28, 43]. This would suggest that prolonged exposure to random errors would lead to a subsequent reduction of uncertainty in the model of the environment reducing the neuromodulatory response, which in turn would reduce the uncertainty in the model of the body. The consequent increase in the reliability of the predictive models of the environment and the body would immediately result in the reestablishment of body ownership. In particular, we expect that after more prolonged exposure to the incongruent stimuli in “I”, the experience of ownership as measured by questionnaires, hand withdrawal, and GSR would return to normal such that there would be no differences in the perceived ownership between the two conditions. Further work shall systematically address this question by running additional trials to assess the temporal evolution of body ownership in the context of an incongruent environment.

What about performance? Our results demonstrate that the sensory manipulations in the incongruent condition did not affect either the self-reported experience of agency (Figure 2) or the performance measured through scores, angular errors (Figure 3) or response times. On the one hand, we designed the paradigm such that all the bodily (i.e., within the peripersonal space), action-driven (reafferent) and task-relevant stimuli were always congruent and therefore fully predictable. Specifically, the 1:1 mapping between the real and the virtual hands ensured that the visual feedback of the movements in virtual reality fully reflected the movements of the real hands resulting in visuomotor congruency. Similar to the peripersonal signals, the consequences of actions in the extrapersonal space signaled by auditory and visual feedback reflected real-world physics and were fully predictable. That is, the sound of the puck temporarily and spatially corresponded to the location of its collision with the environment (i.e., the walls or the goal). On the other hand, we also experimentally controlled for the occurrence of the sensory manipulations to ensure that they are action-independent. Specifically, they were always introduced randomly between the end of a trial and the beginning of the next one. Finally, neither did the manipulated signals in “I” inform about the outcomes of the task (i.e., knowledge of performance) nor did they affect motor performance, which made them task-irrelevant. Indeed, the goal of every experimental session was to complete the motor task as accurately as possible by hitting the puck into the goal. We thus speculate that the congruency (and therefore the predictability) of all the sensory information relevant to the effector and the target in both conditions resulted in an unbiased performance and reinforced the experience of agency, that is the experience of controlling one’s actions, and, through them, events in the outside world [36, 73].

In conclusion, our results support the notion that the plasticity of body ownership depends on an active interplay between the experience-driven top-down predictions and bottom-up prediction errors driven by purely external and action-independent cues which pertain to the environment. Hence, these findings extend current accounts by demonstrating that the sensory evidence necessary for constructing ownership goes beyond the *body* and the peripersonal space [48]. In line with the motor control and perception studies, our data support a functional coupling between the predictive (generative) models of the body and environment [51]. Moreover, our results are consistent with previous findings which demonstrate that body ownership, in fact, affects the perception of certain aspects of the environment (i.e., size of objects) [71], which suggests a bidirectional link between the internal models of the environment and the body. Future work should include a systematic study of the weighting of specific exafferent and reafferent unimodal and multisensory information in modulating the experience of body ownership under different tasks as well as their neural underpinnings.

At this point, however, we expect that the current findings will allow for the advancement of our understanding of the principles underlying the emergence and experience of body ownership, which we propose can be understood in a framework of active inference of all the signals within and outside of the peripersonal space. We believe that the reported results can also contribute to the development of robust computer-based paradigms for the treatment of neurological disorders of cognitive, perceptual, and motor functions.

## Data availability

Data available upon request.

## Authors’ Contributions

K.G., J.T., and B.R.B. designed the protocol; J.T. conceived and conducted the experiment; K.G. and J.T. analyzed the results; K.G., J.T., B.R.B., and P.F.M.J.V. wrote the manuscript. P.F.M.J.V. initiated and supervised the research. All authors reviewed and approved the manuscript.

## Research Ethics

The reported experimental procedures with healthy human subjects followed written consents and were in accordance with the established ethical standards, guidelines and regulations. The study was approved by the ethics committee of the University of Pompeu Fabra (Barcelona, Spain).

